# Activation of nuclear receptors correlates with tuberculosis severity and is a target for host-directed therapy

**DOI:** 10.1101/2025.07.23.666296

**Authors:** Ana Raquel Maceiras, Marta L. Silva, Joana Couto, Rute Gonçalves, Marco Silva, Salvador Macedo, Diana Machado, Iaia Indafa, Armando Sifna, Cesaltina D. Malaca, Nelson I. Namara, Lilica Sanca, Pedro N. S. Rodrigues, Miguel Viveiros, Frauke Rudolf, Christian Wejse, Baltazar Cá, Margarida Saraiva

**Affiliations:** i3S - Instituto de Investigação e Inovação em Saúde, University of Porto, Porto, Portugal; IBMC - Instituto de Biologia Molecular e Celular, University of Porto, Porto, Portugal; Wellcome Sanger Institute, Wellcome Genome Campus, Hinxton, Cambridge, UK; Doctoral Program in Molecular and Cell Biology, ICBAS -Instituto de Ciências Biomédicas Abel Salazar, University of Porto, Porto, Portugal; Global Health and Tropical Medicine, Associate Laboratory in Translation and Innovation Towards Global Health, Instituto de Higiene e Medicina Tropical, Universidade Nova de Lisboa, Lisbon, Portugal; INASA - Instituto Nacional de Saúde Pública da Guiné-Bissau; Bandim Health Project, Indepth Network, Bissau, Guinea-Bissau; Dept of Infectious Diseases, Aarhus University Hospital, Denmark; GloHAU Center for Global Health, Aarhus University, Denmark

## Abstract

The immune response to *Mycobacterium tuberculosis* is accompanied by metabolic adaptations that fuel host immunity, but that are exploited by the pathogen to ensure persistence and growth. Nuclear receptors, such as liver-X-receptors (LXR), orchestrate macrophage immunometabolic adaptations to infection and globally associate with tuberculosis (TB) protection. Here, we show that the “signalling by nuclear receptors” (SNR) pathway is detected in the whole blood of TB patients and that its expression correlates with disease severity. Accordingly, we also show that the activation of the LXR pathway progressively increases in the lungs of *M. tuberculosis*-infected C57BL/6 and C3HeB/FeJ mice. Pharmacologic activation of LXR, specifically at the chronic stage of infection, improved infection outcomes and significantly prolonged the survival of the highly susceptible C3HeB/FeJ mice. Common to both mouse models and to in vitro macrophage infections, LXR activation enhanced bacterial control together with an increase in extracellular cholesterol levels. We propose that progressive LXR activation is required to fine-tune host cholesterol availability during *M. tuberculosis* infections and restrict access to this nutrient during chronic stages of infections. Collectively, we identify the SNR pathway as a potential biomarker of TB severity and timely LXR activation as a candidate host-directed therapy.

## Introduction

Tuberculosis (TB) remains a leading cause of death worldwide. In 2023, an estimated 10.4 million people fell ill with TB and 1.25 million died of this disease [1]. The outcomes of infection by *Mycobacterium tuberculosis* vary from elimination, to latency, to incipient, subclinical or active disease of different severities [2]. These different outcomes are largely determined by the host immune response [3, 4]. The study of the whole blood transcriptome of TB patients revealed a neutrophil-driven type I interferon (IFN)-dependent signature [5], and has provided important cues on host protective versus detrimental pathways [6]. This whole blood signature is recapitulated in *M. tuberculosis* HN878-infected C3HeB/FeJ mice [7], where type I IFN blockade or neutrophil depletion resulted in disease improvement [8]. Thus, pathways discovered in human disease and tested in tractable mouse models hold great potential to advance our understanding of TB pathogenesis.

Macrophages are central to TB pathogenesis, serving as host cells for *M. tuberculosis*, initiating the immune response and adapting to the infection through a series of immunometabolic changes [9], [10]. In vitro, in response to live *M. tuberculosis* infection, macrophages shift their metabolism towards glycolysis [11–14], and display a prominent metabolic signature of cholesterol metabolism [15]. The lungs of *M. tuberculosis*-infected mice also change their energetic metabolic profile [16], with alveolar macrophages being characterized by a strong ox-phos signature, and interstitial monocytes by a glycolytic shift [17–19]. A realignment of the host lipid metabolism also occurs in the granuloma, with a high abundance of cholesterol, cholesteryl esters, triacylglycerols and lactosylceramide identified in the human caseum [20]. This realignment is explored by *M. tuberculosis*, which requires cholesterol for survival during infection [21, 22]. Data from the mouse model of infection show that the cholesterol catabolic pathway of *M. tuberculosis* is implicated in TB pathogenesis, mostly during the chronic phase of infection [21, 23–26] and a transcriptional signature of cholesterol catabolism was found in *M. tuberculosis* isolated from sputum samples of TB patients [27, 28]. Cholestenone, a mycobacterial-derived oxidized derivative of cholesterol, was also detected in the sputum of TB patients, highlighting its potential interest as a novel biomarker in TB [15]. In line with the key role of cholesterol during TB, small chemical compounds capable of inhibiting cholesterol utilization by *M. tuberculosis* have been identified as promising therapeutic tools [29–31]. In particular, perturbing cAMP signaling in the mouse model of *M. tuberculosis* infection impacts disease pathology, bacterial physiology and host responses [32]. Targeting the host cholesterol metabolism also showed promising results in experimental *M. tuberculosis* infection. Administration of all trans-retinoic acid (ATRA), which induces cholesterol efflux in macrophages [33] or of cholesterol-lowering drugs (statins) [34, 35] reduced the bacterial burdens in infected mice.

The nuclear receptor family members, liver-X-receptors (LXR), are key regulators of both intracellular cholesterol homeostasis and inflammation [36, 37], playing a role in protection to several intracellular bacteria [38–40], including *M. tuberculosis* [41]. Single nucleotide polymorphisms in LXR genes associated with genetic susceptibility to TB in the Chinese Han population [42]. Activation of the LXR in cultures of *M. tuberculosis*-infected THP1 cells decreased lipid body formation and improved bacterial control [43, 44]. In contrast, silencing LXRα in THP1 cells led to increased bacterial burdens [45]. In the mouse model of *M. tuberculosis* intra-tracheal infection, LXR deficiency associated with poorer outcomes of disease, whilst their activation improved bacterial control [46]. Given the high homology between human and mouse LXRα and LXRβ, the high expression of LXRα in myeloid cells and the promising findings in the context of infectious diseases and TB, we sought to investigate whether the activation of nuclear receptors is detected in TB patients and whether their in vivo modulation in mouse models that recapitulate the human whole blood transcriptional signature of TB [7] may impact disease outcomes.

Here, we show that the ‘signalling by nuclear receptors’ (SNR) pathway [47] is detected in the whole blood of TB patients across independent TB datasets, and accompanied TB severity. Focusing in the LXR component of the nuclear receptor superfamily, we show its dynamic activation in the lungs of both C57BL/6 and C3HeB/FeJ mice upon aerosol infection with *M. tuberculosis* HN878. Importantly, pharmacologic modulation of LXR improved outcomes of infection in both mouse models, being time-dependent, maximal upon LXR activation in the chronic stage of infection, and connected to cholesterol efflux, without major alterations to the immune response. In all, by combining whole blood transcriptomics of TB patients and experimental mouse data, we reveal a candidate molecular marker of TB severity and demonstrate the potential of timely reinforcing protective pathways, such as LXR-mediated cholesterol efflux, to improve TB outcomes.

## Results

### The signalling by nuclear receptors pathway is activated during TB and correlates with disease severity

To investigate whether the activation of the SNR pathway may be detected in the whole blood of TB patients, we performed gene set enrichment analysis (GSEA) in two previously published RNAseq datasets. The London dataset (referred to as Berry - London) comprises healthy donors, individuals with latent TB and those with active TB, and the South Africa dataset (referred to as Berry - South Africa) includes participants with latent or active TB [5, 48]. The SNR pathway was detected in both datasets, in position 101/ 227 for the Berry London dataset, and in position 112/246 in the case of the South Africa dataset, upon ranking by adjusted p-value (Supp. Table 1 and 2). Of the 315 genes forming the SNR reactome pathway “R-HSA-9006931; signal by nuclear receptors” a total of 25 genes were found to be differentially expressed in at least one of the tested cohorts, being notoriously higher in TB patients as compared with healthy donors or latently infected individuals (Fig. 1A). Since a certain degree of heterogeneity in the expression of the SNR genes was detected at the patient level, we questioned whether this heterogeneous expression might reflect TB disease severity. Using the Berry – London dataset, we analysed how the eigengene expression of the SNR varied according to the extent of radiologic disease based on prior chest x-ray stratification [5]. This analysis was not possible for the South Africa dataset, due to absence of clinical data. Increased expression of the SNR pathway indeed accompanied disease severity, presenting a positive and significant Spearman correlation (ρ = 0.62, p = 0.002; Fig. 1B).

**Figure 1.**
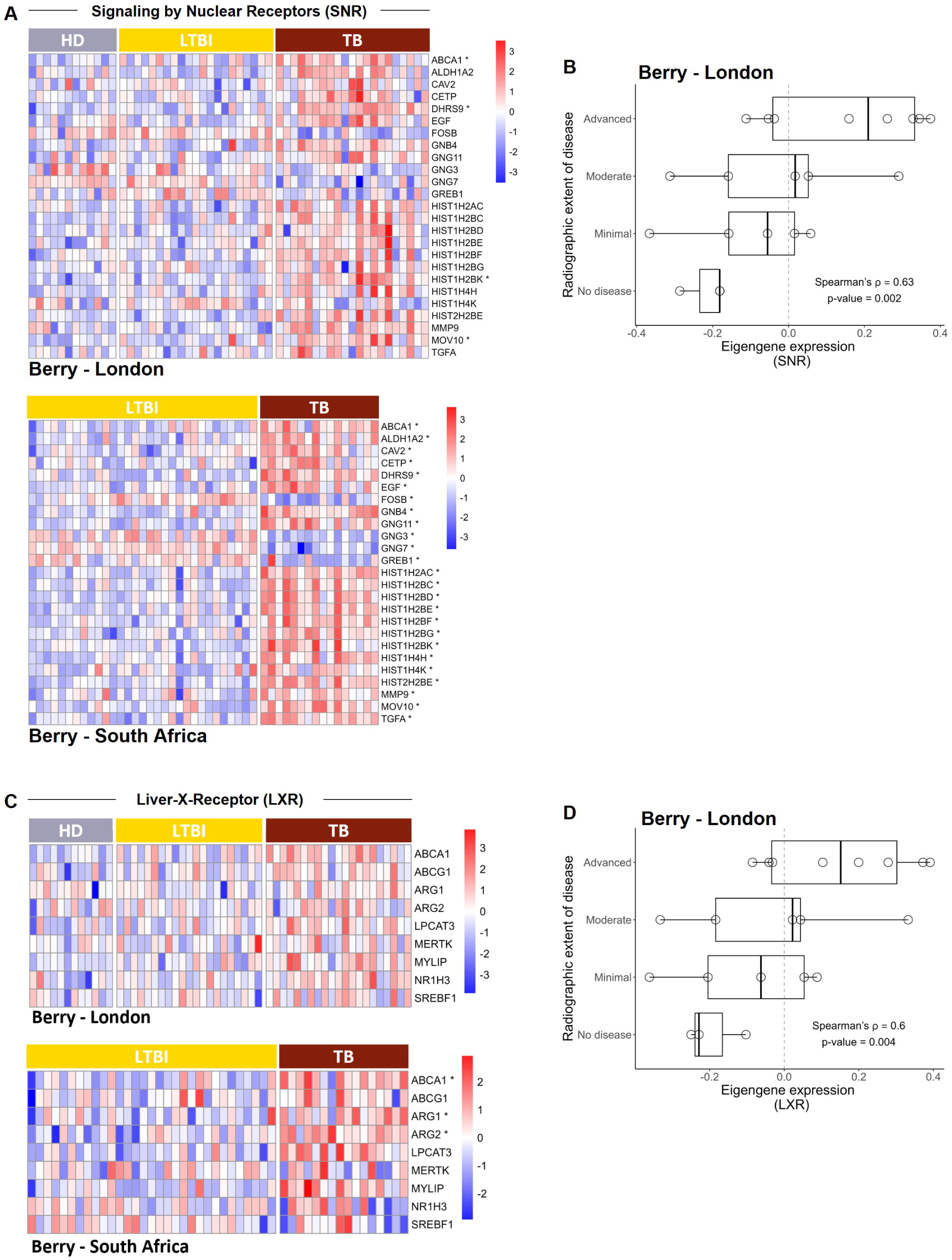
The signaling by nuclear receptors and liver-X-receptor pathways are detected in the blood of TB patients and correlate with TB severity. (A) Heatmaps showing the relative expression of genes detected from the SNR pathway for the Berry-London and the Berry-South Africa cohorts for healthy donors (HD), latent TB infection (LTBI) and TB patients (TB). (B) Violin plot showing the distribution of the eigengene expression of the SNR across the radiographic extent of disease, classifying the TB patients from the Berry-London cohort. (C) Heatmaps showing the relative expression of genes detected from the LXR pathway for healthy donors (HD), latent TB infection (LTBI) and TB patients (TB) from the Berry-London and Berry-South Africa datasets. (D) Violin plot showing the distribution of the eigengene expression of the LXR across the radiographic extent of disease, classifying the TB patients from the Berry-London cohort. In A and C, each column represents an individual; differentially expressed genes for the specific cohort are identified with an *. In B and D, the correlation between the two variables was determined by Spearman correlation analysis, with the ρ and p values indicated.

### Increased expression of the LXR pathway in humans and mice upon *M. tuberculosis* infection

Among the various SNR family members, we next decided to focus our study on the LXR pathway, a member of the nuclear receptor superfamily previously associated with intracellular cholesterol homeostasis and inflammation [36, 37] and TB protection [42–44, 46], two observations that may be linked. As above we interrogated the two whole blood transcriptome datasets, this time for genes that code proteins downstream the LXR receptors involved in cholesterol regulation or inflammatory responses. As for the SNR pathway, the expression of the selected genes of the LXR pathway was heterogeneous and higher in TB patients than in controls (Fig. 1C). Furthermore, we also found a significant positive Spearman correlation (ρ = 0.60, p=0.004) between the LXR pathway eigengene expression and disease severity (Fig. 1D).

We also analysed the activation of the LXR pathway in C57BL/6 and C3HeB/FeJ mice infected via aerosol with *M. tuberculosis* isolate HN878, using previously generated RNAseq datasets for these models at the peak of disease [7]. We found that the LXR pathway was highly represented both in blood (Fig. 2A) and lung (Fig. 2B) of infected mice, regardless of the mouse genetic background. To investigate the kinetics of expression of the LXR pathway at the site of infection, we measured the mRNA transcripts of several genes downstream of LXR activation in the lungs of either mouse strain before infection (day 0) or at days 6, 12 and 25 post-infection. These time points were chosen based on the dynamics of bacterial load accumulation in the lungs of *M. tuberculosis* HN878-infected C57BL/6 or C3HeB/FeJ mice (Supp. Fig. 1A). With the exception of the *Apoe* gene in C57BL/6 mice, whose expression was not altered with infection, the other tested genes were generally upregulated over-time upon infection in both mouse strains (Fig. 2C). These data further support the link between the activation of nuclear receptors and TB disease progression. However, some differences were detected for the expression profile of some genes in C3HeB/FeJ mice between the RNAseq data (Fig. 2A,B) and the lung qPCR (Fig. 2C), which might be related to differences in the initial infection dose, which was high for the RNAseq experiments and low for the qPCR ones.

**Figure 2.**
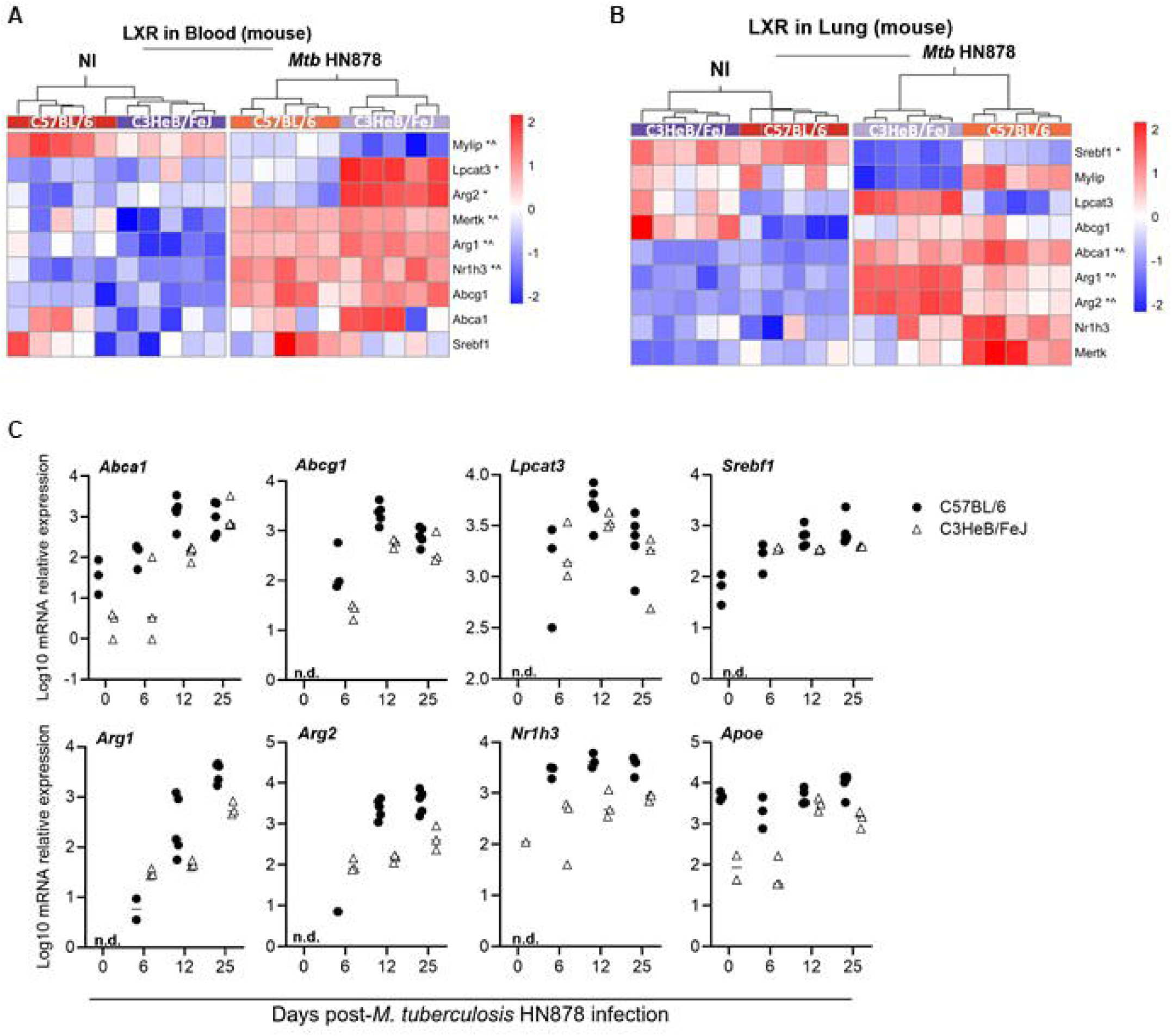
Liver-X-receptor activation in experimental (mouse) TB. (A,B) Heatmaps showing the relative expression of selected genes from the LXR pathway in the blood (A) and lung (B) of non-infected (NI) or aerosol-infected C3HeB/FeJ or C57BL/6 mice. Data are from infections with high doses of *M. tuberculosis* HN878, from a publicly available RNAseq dataset. For both heatmaps, each column represents an individual mouse, normalized gene expression by gene (Z-score by row) is presented and differentially expressed genes are identified with an *, for C3HeB/FeJ mice or ^, for C57BL/6 mice. (C) Transcriptional analysis of the indicated genes of the LXR pathway by qPCR in the lungs of C57BL/6 (circles) or C3HeB/FeJ (triangles) mice aerosol-infected with *M. tuberculosis* isolate HN878 (580 and 209 CFU delivered to the lung of C57BL/6 or C3HeB/FeJ mice respectively), prior to (day 0) or at different time-points (day 6-12-25) post-infection. Each dot represents an individual mouse in one experiment. n.d., not detected

### The beneficial effect of LXR activation during *M. tuberculosis* aerosol infection of C57BL/6 mice is time-dependent

LXRαβ double-deficient mice were shown to be more susceptible to intra-tracheal *M. tuberculosis* infection, displaying higher CFUs and lung histopathology, whereas activating the LXR pathway prior or post-infection demonstrated improved lung bacterial burdens, possibly due to increased early neutrophil recruitment and enhanced CD4 T cell responses [46]. However, the pharmacologic activation of this pathway during aerosol infection of mouse models that better recapitulate the human whole blood TB signature had not been tested. Furthermore, considering our findings showing a dynamic transcription of the LXR pathway during *M. tuberculosis* infection, we hypothesised that the timing of LXR potentiation might impact the final result. To investigate these hypotheses, we administered the LXR agonist T0901317 to C57BL/6 mice 7 days prior to infection, or day 6, 12 or 18 post-*M. tuberculosis* HN878 aerosol infection. Prophylactic administration of the LXR agonist or its administration on days 6 or 12 post-infection did not impact the weight loss curve of vehicle or LXR-activated infected mice (Fig. 3A, top panels). In contrast, mice that received the LXR agonist from day 18 post-infection did not lose as much weight as the vehicle group (Fig. 3A, top panels), suggesting a time-dependent beneficial effect of the administered regimen. On day 24/25 post-infection, at which point the control (vehicle) group reached the peak of disease, we analysed the lung bacterial burdens for all groups. Prophylactic administration of the LXR agonist resulted in a minimal decrease of lung bacterial burdens (Fig. 3A, bottom panel). Early activation of the LXR pathway (day 6 post-infection) did not impact lung bacterial burdens, whereas activation from day 12 post-infection resulted in a modest decrease in lung bacterial burdens (Fig. 3A, bottom panel). Strikingly, a pronounced decrease of over a log10 in bacterial numbers in the lungs of treated mice was observed upon activation of the LXR pathway from day 18 post-infection (Fig. 3A, bottom panel).

**Figure 3.**
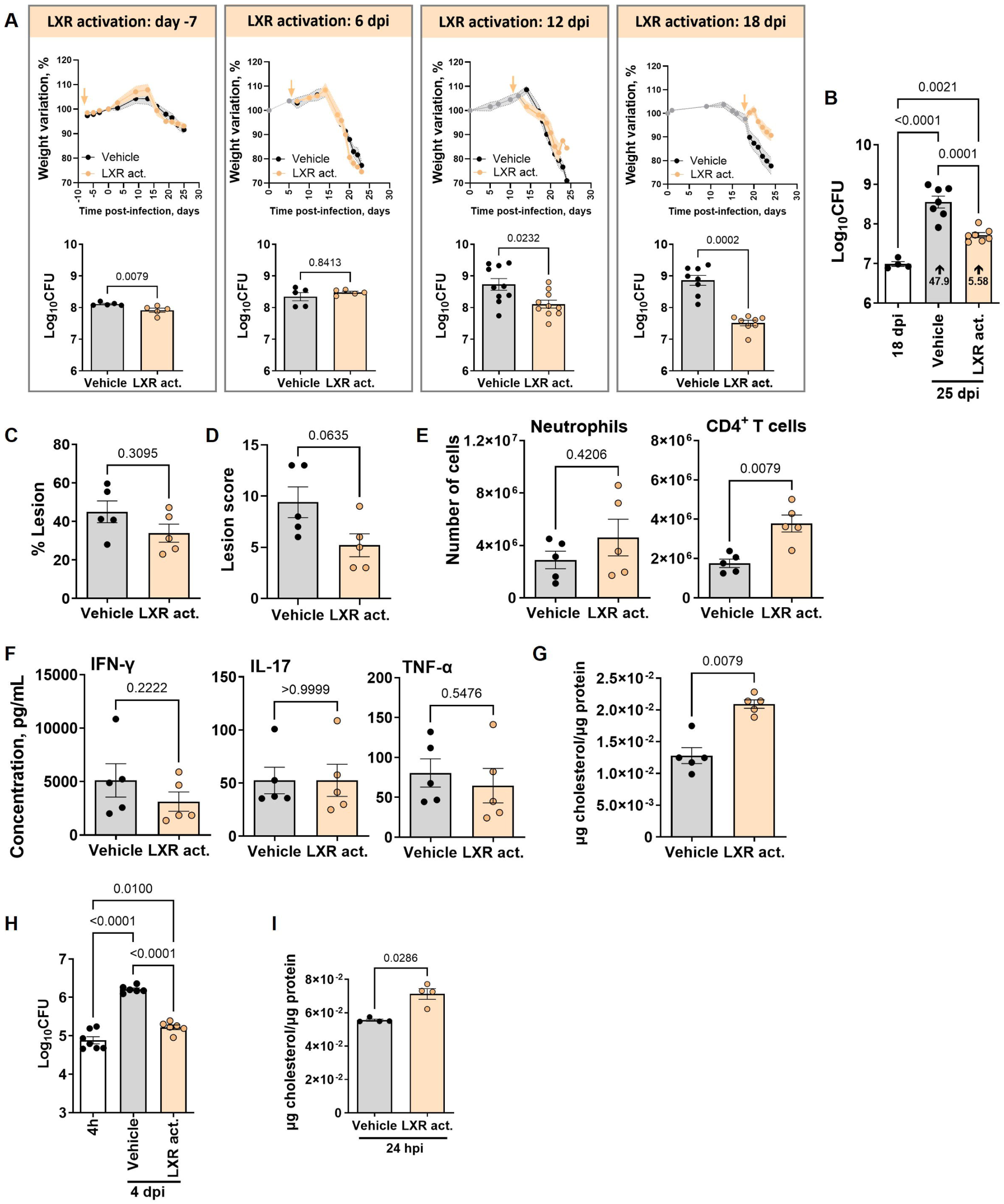
Time-dependent effect of liver-X-receptor activation in controlling *M. tuberculosis* infection of C57BL/6 mice. C57BL/6 mice were treated with the LXR activator T0901317 at days 7 prior to infection (−7), or days 6, 12 or 18 post-aerosol infection with *M. tuberculosis* isolate HN878 (with infections ranging from 503 to 853 CFU delivered to the lung). (A) Body weight variation for control (vehicle; grey) or treated (orange) mice during the course of the experiment (top panels) and bacterial burdens in CFU determined on day 25 post-infection (bottom panels). The arrows in the top panels represent the initial time-point of LXR agonist administration. (B) C57BL/6 were infected as above and bacterial loads determined on days 18 (prior to vehicle or LXR administration) and 25 (experimental end-point) post-infection. The fold increase in lung bacterial burden from day 18 to day 25 in each experimental group is represented in the respective bars. (C) For the mice infected in (A) and treated with the LXR activator from day 18 post-infection, the % of lung lesioned area (C), lesion score (D), number of neutrophils and CD4 T cells in the lung (E), levels of the indicated cytokines (F) and total cholesterol (G) in the lung supernatants were determined on day 25 post-infection for control (vehicle; grey bars) or treated (orange bars) mice. Each dot represents the mean of five animals with SEM shaded (A, top panels) or independent mice (mean ± SEM; A, bottom panels, B-G). Activation of the LXR pathway at - 7 or +6 days was tested once, and on days 12 or 18 in two independent experiments. (H, I) Bone marrow-derived macrophages (BMDM) from C57BL/6 mice were infected with *M. tuberculosis* HN878 (MOI=2) and 4 h later treated or not with 100 nM of the LXR activator. Bacterial loads (H) were determined 4 h and 4 days post-infection and total cholesterol detected in the supernatants of the infected cultures 24 h post-infection (I). Data are mean ± SEM, with each dot representing one individual well. Mann-Whitney U tests were used to identify statistical differences between the two groups. p-values are indicated and considered significant if ≤ 0.05. dpi, days post infection; hpi, hour post-infection; Mtb, *M. tuberculosis*.

Focusing on the effect of LXR administration at this time-point, we investigated whether the observed protection might be due to restriction of bacterial growth. The increase in the lung bacterial burden from day 18 to the end-point of the experiment (day 25) was much larger in control mice than in those that received the LXR agonist (respectively, 47.9 versus 5.58 increase; Fig. 3B). Despite this improved control of bacterial growth, the % of lesioned lung area (Fig. 3C) was not significantly altered in LXR-treated mice and the lung lesion inflammatory score showed only a modest decrease (Fig. 3D; Supp. Fig. 1B). In an apparent paradox with its increased expression as the infection progresses, these data suggest that reinforcing the LXR pathway near the peak of infection improved bacterial control with no major effect on tissue damage restriction.

To further investigate this apparent paradox, we next analysed the immune cell populations present in the lungs of vehicle or LXR-treated mice (Supp. Fig. 1C). An overall increase of the various immune cell populations was detected in LXR-treated mice (Supp. Fig. 1D). Of particular interest in determining pathology versus protection in TB are neutrophils versus CD4 T cells [7]. Notably, *M. tuberculosis*-infected mice that received the LXR agonist on day 18 post-infection showed similar numbers of neutrophils and increased numbers of CD4 T cells (Fig. 3E), as compared to control mice. The general increase in lung immune cells, together with a similar presence of neutrophils, in LXR-treated mice may explain the fact that the lung infiltrated area and histological score remain similar to that of control mice, despite improved control of bacterial growth. Because the increase in the numbers of CD4 T cells was not accompanied by an increase of effector cytokines namely IFN-γ, IL-17 or TNF in the lung supernatants (Fig. 3F), nor by a different organization of lung lesions in the lungs of LXR-treated animals (Fig. 3C, D), we hypothesised that the protective effect of LXR activation may be linked to other mechanisms.

### LXR activation enhances the macrophage bacterial control and the extracellular levels of cholesterol

Given the relevance of the LXR pathway to cholesterol metabolism [49] and of cholesterol to *M. tuberculosis* pathogenesis [50], we hypothesised that pharmacological activation of the LXR may enhance cholesterol efflux from host cells depriving *M. tuberculosis* of this carbon source. This would be in line with the enhanced expression of cholesterol transporters induced by the LXR agonist T0901317 in *M. tuberculosis*-infected THP1 cells [43]. Indeed, we detected increased levels of cholesterol in the supernatants of lungs of *M. tuberculosis* HN878 infected C57BL/6 mice treated with the agonist from day 18 to day 25 post-infection (Fig. 3G). To investigate if this effect of LXR activation on the host was time-dependent, we activated the LXR pathway between days 6 and 12. LXR activation during this period did not impact the levels of extracellular cholesterol (Supp. Fig. 2A). Also, at the end of the LXR activation, only a minor effect was observed in bacterial burdens (Supp. Fig. 2B), which was not sustained over time, as by day 25 post-infection no differences in the lung CFUs were observed between the two groups (Supp. Fig. 2C).

Since previous studies demonstrated that LXR activation in THP1 cells limited *M. tuberculosis* growth [43, 44], we next infected mouse bone marrow-derived macrophages (BMDM) with *M. tuberculosis* HN878 and allowed the infection to progress for 4 days in the absence (vehicle) or presence of the LXR activator. Activation of the LXR pathway decreased intracellular bacterial growth as compared to vehicle controls (Fig. 3H), while increasing the levels of extracellular cholesterol (Fig. 3I). This protective effect of the LXR activator in BMDM was not exclusive to infection with *M. tuberculosis* HN878, as similar results were obtained upon infection of macrophages with a clinical isolate from lineage 4 (Supp. Fig. 2D). Comparable findings were observed in THP1 cells (Sup. Fig. 2E). In line with our in vivo data, increased bacterial control was invariably accompanied by increased extracellular cholesterol amounts (Sup. Fig. 2D, E). Since activation of the LXR pathway in macrophages reduced *Salmonella enterica* serovar Typhimurium phagocytosis [39], we tested if a similar mechanism would operate in *M. tuberculosis*. However, no differences in bacterial burdens were detected in the presence of the LXR agonist during the first 4 h of infection (Supp. Fig. 2F). Furthermore, the presence of the LXR activator in axenic media did not directly impact the growth of *M. tuberculosis* even when cholesterol was provided as an energy source (Supp. Fig. 2G), which supports a host-mediated effect of the LXR activator.

### Impact of LXR activation in *M. tuberculosis* infections of C3HeB/FeJ mice

Having demonstrated a protective effect of late LXR activation in C57BL/6 mice, we tested the same intervention in C3HeB/FeJ mice, which better recapitulate the immune and histological features of human TB [7, 8]. Treatment of C3HeB/FeJ mice with the LXR activator at 18 days post-aerosol infection with *M. tuberculosis* HN878 led to a decrease in lung bacterial burdens (Fig. 4A) and an increase in the levels of extracellular cholesterol (Fig. 4B). The differences seen between vehicle and LXR-activated C3HeB/FeJ infected mice recapitulated those detected for C57BL/6 ones, but were less pronounced. In contrast with the general increase in immune cells observed in the case of LXR-treated C57BL/6 mice (Supp. Fig. 1D), activation of LXR in infected C3HeB/FeJ mice did not change most of the lung immune cell populations (Supp. Fig. 3A). However, also in sharp contrast with what was observed in C57BL/6 mice, it resulted in a significant increase in the number of lung neutrophils and a significant decrease in CD4 T cells (Supp. Fig. 3A).

**Figure 4.**
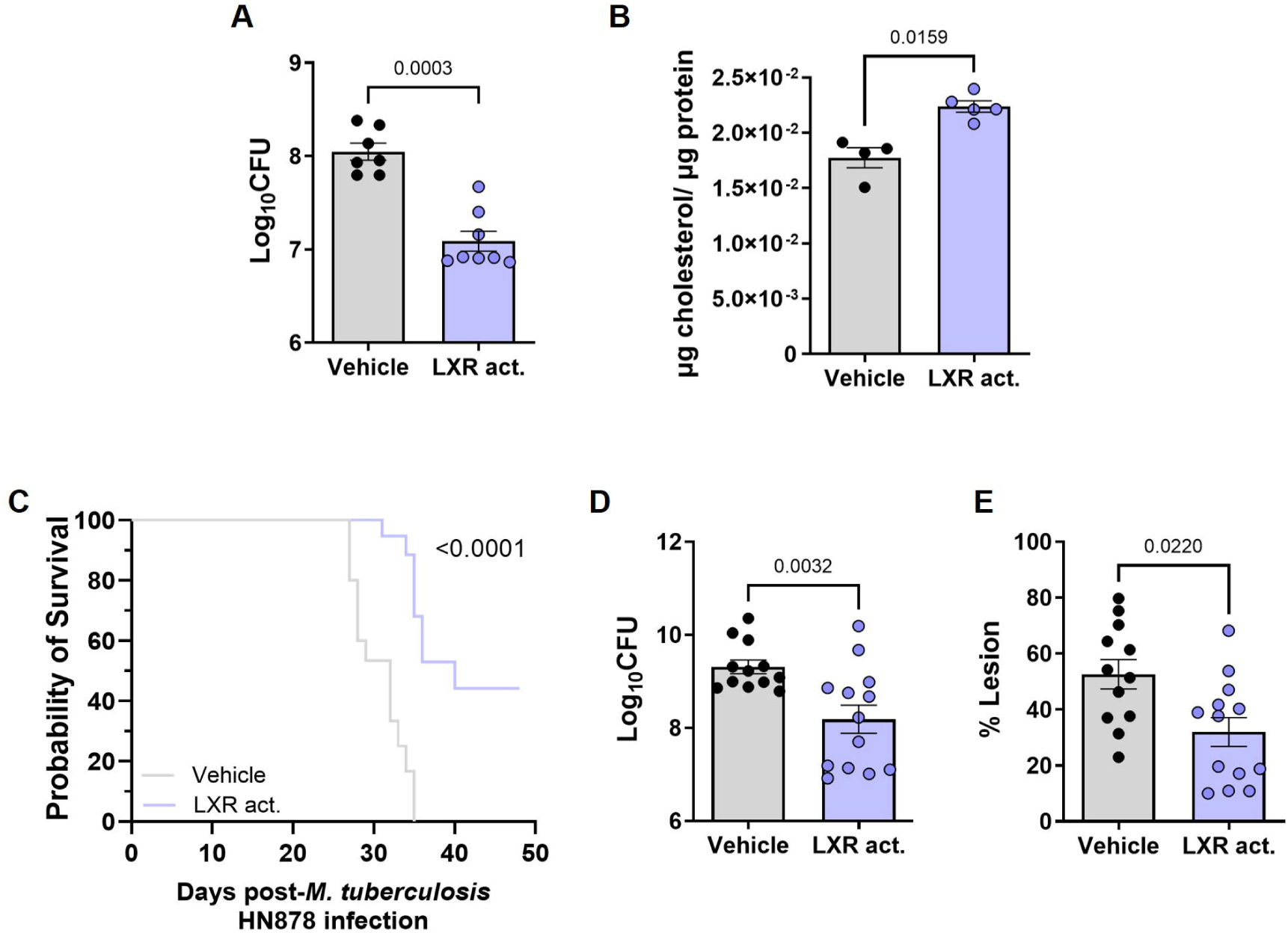
LXR activation reduces lung bacterial burdens and prolongs survival of *M. tuberculosis*-infected C3HeB/FeJ mice. C3HeB/FeJ mice were aerosol-infected with *M. tuberculosis* isolate HN878 (203 or 271 CFU delivered to the lung) and administered with the LXR activator T0901317 or vehicle control from day 18 post-infection. On day 24 post-infection, the lung bacterial burdens (A) and total cholesterol in the lung homogenates (B) were quantified. (C) Survival curve (humane end-points defined based on over 20% weight loss or severely compromised respiratory function) for C3HeB/FeJ animals infected with *M. tuberculosis* as above and treated with vehicle or the LXR agonist for up to 50 days. Lung bacterial burdens (D) and percentage of lung lesioned area (E) for each animal enrolling the survival curve at its respective end-point. (A, B, D, E) Vehicle and treated groups are represented by grey or blue bars, respectively. Data are mean ± SEM. Each dot represents an individual mouse. (C, D, E) Data is pooled from 2 independent experiments (dose of infection 203 and 271). Mann-Whitney U tests were used to identify statistical differences between the two groups. p-values are indicated and considered significant if ≤ 0.05. In (C) A Log-rank (Mantel-Cox) test was used to compare the survival curves; p-values are indicated and considered significant if ≤ 0.05.

An increase in neutrophils and decrease in CD4 T cells has been associated with TB susceptibility in different studies (ref). Thus, we questioned whether prolonging the LXR activation in infected C3HeB/FeJ mice would indeed improve disease outcomes or only transiently enhance bacterial control. To test this, the administration of the LXR activator was prolonged up to day 50 post-infection. Whereas the vehicle-treated animals showed pronounced weight loss (Supp. Fig. 3B), reaching humane end-points before day 40 post-infection, those receiving the LXR activator survived significantly longer, with 6/12 animals still performing well at the end-point (Fig. 4C; Supp. Fig. 3B). Of note, when pooling all animals irrespective of their time-of-death, the LXR-treated C3HeB/FeJ mice showed lower bacterial burdens and lesioned area than vehicle controls (Fig. 4D, E). Therefore, LXR administration also improved the outcome of infection in C3HeB/FeJ mice, significatively prolonging their survival.

## Discussion

TB remains a high-burden disease, with important challenges being the development of improved diagnosis methods and better, less toxic and shorter regimens [51]. The study of the whole blood transcriptome of TB patients has allowed the identification of immune signatures of disease, together with their phenotypic heterogeneity, evolution during infection and resolution upon treatment [5, 48, 52]. Moreover, it has shed important light into the mechanisms of TB pathogenesis advancing the discovery of potential targets for host-directed therapies [6, 8]. Here, we resorted to available transcriptomic datasets of TB patients and *M. tuberculosis*-infected mice to investigate the expression of nuclear receptors during TB. Nuclear receptors are a superfamily of transcriptional regulators linking lipid metabolism, inflammation and the immune system [47], with a relatively well established role in TB [41]. We show that the overall SNR pathway was enriched in TB patients from London and South Africa, as compared to the respective healthy controls. Moreover, for the London dataset, we found that the extent of SNR activation in the blood increased with the radiologic extent of disease. Thus, our study provides initial evidence of the SNR pathway as a molecular candidate of TB disease progression and, potentially, TB severity. Even though our findings are validated in different sets of patients, the small numbers included represent a limitation of the study. Future validation of the SNR as a relevant marker in TB is important, as stratifying TB patients is increasingly acknowledged as a key step to devise rational strategies to treat TB, namely to adjust the required duration of current regimens in a patient-centred way [51].

We further focused our study on the LXR pathway, a specific molecular component of the SNR, because LXRs are expressed in macrophages, the preferred cellular niche for *M. tuberculosis*, and their activation regulates the metabolism of cholesterol, an important nutrient for this pathogen across different phases of infection, from phagocytosis to chronicity [53]. Furthermore, previous studies highlight the protective effect of LXRs during in vitro [43–45] and in vivo [46] *M. tuberculosis* infections. We detected the activation of selected genes of the LXR pathway in the blood of TB patients across the aforementioned datasets, and reported a positive correlation between the LXR eigengene and the extent of radiologic disease, further hinting at the relevance of this pathway in TB. Moreover, we investigated the dynamics, effects and underlying mechanisms of the LXR activation in mouse models that reflect key features of the human disease: aerosol route of infection with an isolate of *M. tuberculosis* that allows for a whole blood transcriptomic signature recapitulating that seen in TB patients [7]. Our study thus departs from a previous one using intra-tracheal infection with *M. tuberculosis* H37Rv laboratory reference strain [46], confirming some findings, but also revealing new ones. We report a progressive increase of the expression of several genes downstream the LXR pathway in the lungs of *M. tuberculosis* HN878-aerosol infected C57BL/6 and C3HeB/FeJ mice, and also detected their expression at the peak of disease in the lungs and blood of these mice. This enhanced lung expression of the LXR pathway as the infection progresses is in line with our human data and with a previous study, which tested a smaller set of LXR responsive genes in the bronchoalveolar lavage cells of mice infected intra-tracheally with *M. tuberculosis* H37Rv strain [46]. It will be important in future studies to characterise the activation of the LXR pathway during infection beyond gene expression, as well as to identify which molecular cues trigger it.

Collectively, our analyses of the SNR/LXR expression in human TB and mouse models support a link between the activation of these pathways and disease progression. In the context of the established protective role of LXR in TB [42–46], these findings may seem counterintuitive: how would a pathway whose expression increases with disease severity be protective? Further adding to this apparent paradox, in our experimental models, reinforcing the LXR pathway prior to or early on post-infection did not result in enhanced protection. The lack of effect of prophylactic LXR activation contrasts previous results [46], possibly due to the difference in the route of administration (aerosol versus intra-tracheal) and the strain of *M. tuberculosis* used (HN878 versus H37Rv). Interestingly, reinforcing the LXR pathway at the peak of its natural activation markedly improved *M. tuberculosis* growth control by the two mouse strains. In particular, this intervention significatively prolonged the survival of the highly susceptible C3HeB/FeJ mice. Thus, the LXR activation by the host is likely to both reflect the progression of infection and represent a mechanism to restrict *M. tuberculosis* growth triggered during chronic stages of infection. The molecular cues activating and regulating LXR activation remain unknown, but it is possible that the accumulation of oxysterols in the lungs of infected mice play a role [54]. This accumulation may result from metabolic alterations in the host cells, notably macrophages, but also be promoted by the pathogen, as *M. tuberculosis* encodes several enzymes capable of hydroxylating cholesterol and generating LXR agonists [25, 55–57]. Oxysterols are gaining importance as immune modulators in bacterial and viral infections [58] and are therefore of interest to further study in the context of TB.

We hypothesised that the mechanism underlying the time-dependent beneficial effect of LXR activation might be linked to alterations to intracellular cholesterol availability, a key nutrient for *M. tuberculosis* specifically during the chronic stage of infection [21, 23, 24, 26, 59]. An increase in the extracellular levels of cholesterol was indeed detected in vivo in the lungs of both C57BL/6 and C3HeB/FeJ mice that benefited from receiving the LXR agonist upon infection. Moreover, a similar increase was detected in vitro in BMDM cultures infected with *M. tuberculosis* HN878, or with an isolate of *M. tuberculosis* belonging to lineage 4. This is relevant, because strain specific differences in the ability to metabolise lipids, specifically cholesterol have been reported [60]. Although it is impossible to capture the full breath of host-*M. tuberculosis* interactions, our findings suggest cholesterol efflux and subsequent nutrient deprivation as the common axis linking the pharmacological activation of LXR with *M. tuberculosis* growth restriction. Of note, the LXR-mediated protection may however be disrupted in the context of TB and metabolic comorbidities, such as diabetes [54]. Also of note, other cholesterol targeting strategies have been tested during *M. tuberculosis* infection [33–35], but none reduced the bacterial burdens in infected mice to the extent we report here. This higher success of the LXR activation may be due to different modes of action, as well as experimental design, namely the time of activation which we show to be an important variable.

Interestingly, our in vivo data suggest that a remodelling of the immune response induced by LXR activation may not be a determinant factor underlying protection. In *M. tuberculosis*-infected C57BL/6 mice, administration of the LXR agonist contributed to an overall increase of immune cells in the lungs, the only exception being neutrophils. This global increase of immune cells may explain the fact that the lung infiltrated area remained similar to that of control mice. The increase of CD4 T cells is in line with a previous report of LXR activation during intra-tracheal *M. tuberculosis* infection [46]. However, unlike that study, we saw no evidence of increased Th1/Th17 cell responses, as the amounts of IFN-γ and IL-17 proteins present in the lung supernatants were comparable in control or LXR-treated mice. In C3HeB/FeJ infected mice, the activation of the LXR pathway resulted in decreased CD4 T cell and increased neutrophils being recruited to the lung. This divergence of protective versus detrimental immune responses in C57BL/6 versus C3HeB/FeJ mice, did not compromise the ultimately similar outcome of *M. tuberculosis* control. Thus, we propose a model where in addition to any possible adjustments to the immune response caused by LXR activation, nutrient limitation may act as the main mechanism for the beneficial action of LXR potentiation in tractable mouse models of TB.

Taken together, our findings suggest that progressive LXR activation may be required to fine-tune macrophage responses and cholesterol availability during *M. tuberculosis* infections. During initial stages of infection low LXR activation might be required to limit its anti-inflammatory function [37] and ensure that cholesterol is available for key macrophage functions [53]. Later on, activation of LXR likely restricts the pathogen’s access to cholesterol, which is a key nutrient for *M. tuberculosis* during chronic stages of infections. It is important to mention that at different host-pathogen interfaces, pharmacological modulation of the LXR pathway may yield similar host-protective results through distinct complex regulatory actions. For example, LXR activation is also protective during S. Typhimurium infections, but in that case involve CD38/NAD/cytoskeleton rearrangements which impact the phagocytic process [39].

In summary, our study expands the growing field of host metabolic regulation during infection, by proposing the SNR pathway as a possible biomarker of TB severity and cholesterol modulation by timely LXR activation as a host-directed therapy of interest in TB. Rational stratification of TB patients may impact the design of better therapeutic strategies, while improving LXR agonists for clinical use in TB, as well as testing their effects at even later time-points post-infection (once the lung lesions show the typical consolidated structure) alone or as adjuvants to TB antibiotherapy, are exciting avenues of research. Finally, our study contributes to two broader concepts in TB: that not all pathways reflecting disease severity might be detrimental, and that reinforcing protective pathways from a therapeutic perspective may require timely interventions.

## Methods

### Study approval

Animal experiments followed the ARRIVE guidelines, the 2010/63/EU Directive and were approved by the i3S Animal Ethics Committee and the Portuguese National Authority for Animal Health (DGAV; #018413/2021-11-24).

### Sex as a biological variable

Our study examined male and female mice, and similar results were found for both sexes.

### RNAseq data analyses

Mouse whole-blood and lung RNAseq data from [7] were downloaded from SRA (GEO accession code GSE140945). The quality of raw sequencing data and the presence of adaptors were initially assessed using FastQC. Low-quality reads with less than 36 nucleotides long, adapters, leading bases with quality below 3, and trailing bases when the average quality per base was below 15 for regions of 4 bases were removed using Trimmomatic v0.39 [61]. HISAT2 v2.2.1 was then used to align the remaining reads to the *Mus musculus* genome Ensembl GRCm38 (release 84) followed by the usage of StringTie v2.1.6 to obtain gene-level counts [62, 63]. The obtained counting table as well as whole-blood RNAseq count matrices from the Berry London and Berry South-Africa cohorts [48] (downloaded from GEO with accession codes GSE107991 and GSE107992, respectively) were imported into R v4.1 for downstream analysis. Raw counts were processed using edgeR v3.36.0 and limma v3.50.0 packages [64, 65]. Initial processing of raw counts was performed with edgeR, and genes with counts per million (CPM) lower than 10 in at least 12 samples were discarded from further analysis. Differential expression analysis on the remaining genes was performed using generalized linear models from the limma package. The effect of the sex covariate was taken into account by adding it as a variable to the model. Genes were deemed significant if log2 fold change was ≥ 1 or ≤ −1 and the false discovery rate (FDR) p-value ≤ 0.05 after Benjamini–Hochberg correction for multiple testing. Plots were obtained using ggplot2 and pheatmap [66, 67]. GSEA was performed on the logFC (aTB vs HD) for Berry London and logFC (aTB vs LTBI) for Berry South Africa, for all genes using ReactomePA [68].

### Bacteria growth and quantification

*M. tuberculosis* HN878 and the L4 clinical isolate were expanded in Middlebrook 7H9 (BD Biosciences, Cat. #90003-876) liquid medium supplemented with 10% OADC and 0.2% glycerol (Sigma-Aldrich, Cat. #G5516-500M) and when in mid-log phase bacterial suspensions were aliquoted in cryovials, frozen, and stored at –80°C. Bacterial quantification was performed by thawing six vials of each stock and plating serial dilutions in 7H11 agar medium supplemented with 10% OADC and 0.5% glycerol. The plates were incubated 21 to 28 days at 37 °C before colony enumeration.

### Animal housing, infection, LXR agonist administration and monitoring

C57BL/6 and C3HeB/FeJ mice were bred and housed at the i3S animal facility. Males and females of 8 to 12 weeks old were used for infections, and maintained under contention conditions in the animal biosafety level 3 facility at i3S, in controlled temperature (20-24°C), humidity (45-65%), and a light cycle of 12h (light/dark). Water and food were provided *ad libitum*. Mice were infected with *M. tuberculosis* (isolate HN878) through the aerosol route using an inhalation exposure system (Glas-Col) [7, 8, 69]. To determine the dose of infection, the bacterial load in the lungs of 3 mice was quantified 3 days post-infection. Doses of infection are indicated in the Figure legends. Infected mice were weighed every week, or every other day when showing signs of disease. Mice were euthanized at the indicated time points or when reaching humane endpoints (weight loss over 20% or poor responsiveness to physical stimulation). For the LXR modulation, mice were treated with 50 µg of LXR agonist T0901317 (Abcam, Cat. #ab142808) or with the vehicle (0,1% DMSO/PBS) administered intraperitoneally, starting on day 7 pre-infection or days 6, 12 or 18 post-infection, every other day until the end of the experiment.

### Lung processing

Lungs were aseptically excised and processed as previously described [7, 8, 69]. Briefly, lungs were digested with Collagenase D (Roche, Cat. #11088882001) followed by physical disruption and filtration through 70 μM cell strainers (Falcon, Cat. #352350.0). Cell suspensions were used for colony forming units (CFU) determination, flow cytometry and mRNA analysis.

### CFU determination

Lung cell suspensions were lysed with 0.1% saponin, serial dilutions prepared and plated in Middlebrook 7H11 agar (BD Biosciences, Cat. #212203.0) supplemented with OADC and PANTA (BD BioSciences, Cat. # 245114). CFU were enumerated after 21-28 days of incubation at 37°C.

### RNA extraction and qPCR

Total RNA was extracted from infected mouse lungs using TRIzol reagent (GRiSP, Cat. #GB23.0100) according to the manufacturer’s instructions. cDNA was synthesized with the SuperScript First-Strand Synthesis System for RT-PCR (ThermoScientific, Cat. #E6300L). Target gene mRNA expression was quantified by real-time PCR, using Taqman Primer Probes specific for the target genes, normalized to *Hprt1/Hmbs* (Taqman; Supp. Table 3).

### Flow cytometry

Mouse lung cell suspensions were stained for surface antigens for 30 min at 4° C and fixed for 20 min in 4% paraformaldehyde-PBS after erythrocyte lysis. Dead cells were excluded using ZombieAqua (Biolegend, Cat. #423102). Cells were acquired on a BD Fortessa II and data was analysed using *FlowJo* software (v10.1.r7). All antibodies used are listed in Supp. Table 4. Gating strategies are shown in Supp. Fig. 1C.

### Histological analysis

Whole lungs were perfused *in situ* with PBS, the right upper lobe was excised, fixed in 10% buffered formalin and embedded in paraffin. Serial 3-μm thick sections were performed and used for hematoxylin and eosin (H&E) staining. Morphometric analysis of the lung pathology was performed as before [69, 70], with the software Interactive Learning and Segmentation Toolkit (Ilastik version 1.3.3). The probability maps of the whole lung and the lesions were analyzed in CellProfiler Analyst (version 3.1.5). Lesion percentage was defined by the area occupied by lesion in each lung and lesion score by scoring histopathological features (peribronchiolitis, perivasculitis, alveolitis and necrosis/cell debris; Supp. Fig. 1B) as previously reported [71].

### Cytokine determination by multiplexed immunoassay

Cytokine levels in the lung supernatants were measured using the LEGENDplex™ Mouse Inflammation Panel kit (Biolegend, Cat. #740446_Mouse Inflammatory Panel) following manufacturer’s specifications. The beads were acquired in an BD Accuri C6 (BD Biosciences), and the data analysed using the LEGENDplex™ Software version 8 (BioLegend).

### In vitro cultures and infections

Mouse BMDMs were generated from C57BL/6 bone marrow suspension cells as previously described [14, 72]. On day 7 of culture, the macrophages were recovered, plated in 24 well-plates at a density of 1×10^6^ cells ml^-1^ and infected with the indicated *M. tuberculosis* isolates at a multiplicity of infection (MOI) of 2. THP1 cells were grown according to ATCC instructions, differentiated with 100 nM PMA (Sigma-Aldrich, Cat. #16561-29-8) for 24 h, followed by a 48 h rest before infection and then plated in 24 well-plates at a density of 1×10^6^ cells ml^-1^ and infected with *M. tuberculosis* HN878 at a MOI of 1. Where indicated, the LXR agonist T0901317 was added to the cultures at a concentration of 100nM. Bacterial burdens were determined at 4 h and 4 days post-infection by CFU enumeration; and the culture supernatants were collected 24h post-infection for cholesterol analysis. Uninfected cells were used as controls.

### Cholesterol quantification

Cholesterol content in the supernatants was measured using the Cholesterol Quantification kit (Sigma-Aldrich, MAK043-1KT) following the manufacturer’s instructions. The amount of cholesterol was normalized to that of total protein detected using a BCA protein Assay kit (Thermofisher, Cat. #23225).

### Growth of *M. tuberculosis* in axenic media

*M. tuberculosis* HN878 was grown in 7H9 supplemented with 10% OADC and 0.05% Tween-80 (Sigma, Cat. #P1754) until mid-exponential phase (OD_600_ of 0.6-0.9). At this stage, a single bacterial suspension was obtained and bacteria were plated in 96-well plates at a final OD of 0.01 in 7H9 medium alone, or with cholesterol and in the presence or absence of the LXR activator T0901317 at a range of concentrations from 0.01 to 10 µM. Cholesterol (Sigma, Cat. #C3045) was dissolved in a solution of tyloxapol and ethanol (Sigma, Cat. #1.00983) (1:1) and added to the medium at a concentration of 200 µM. Cultures were replenished with cholesterol at day 7. Each condition was tested in triplicate. Growth was monitored by measuring the optical density (OD_600_) over the time course of the experiment.

### Statistical analysis

Data were analysed using GraphPad Prism software v8.1.0 or R v4.1. Spearman correlation analysis was performed between the "Radiographic extent of disease" and the SNR and LXR pathways activation, for the Berry -London cohort, to determine the existence of a significant correlation between the two variables. Normality and log normality were assessed using Shapiro-Wilk and outliers with Grubbs test. Statistically significant differences between two groups were determined using non-parametric two-tailed Mann–Whitney U tests. A Log-rank (Mantel-Cox) test was used to analyse the survival curve. Differences were considered significant for p ≤ 0.05.

## Data availability

Data are available from the corresponding author upon request.

## Author contributions

Conceptualization: ARM, MLS, CW, BC, MS

Formal analysis: ARM, MLS, BC, MS

Funding Acquisition: MS

Investigation: ARM, MLS, JC, RG, SM, DM, II, AS, FR, BC, MS

Project administration: FR, BC, MS

Resources: LS, DM, MV

Supervision: PNSR, FR, CW, BC, MS

Visualisation: ARM, MLS, JC, RG, MS

Writing of the draft: ARM, MLS, MS

Writing-review and editing: all authors

Shared authorship: the first two authors share authorship based on the equal amount of work they contributed with. The order was decided based on the fact that ARM initiated the study and provided laboratory supervision to MLS.

## Acknowledgements

This work was funded by National Funds through FCT—Fundação para a Ciência e a Tecnologia, I.P., under the project UID/4293/2025, and by the La Caixa Foundation, grant HR21-00415 to MS. MLS and RG are funded by FCT PhD scholarships 2020.05061.BD and 2022.12852.BD. FCT also finances GHTM (UID/Multi/04413/2020) and LA-REAL (LA/P/0117/2020). DM is funded by FCT through *Estímulo Individual ao Emprego Científico* (CEECIND/00241/2017/CP1386/CT0002). We acknowledge the i3S scientific platforms Animal House, Translational Flow Cytometry, Genomics and Histology and Electron Microscopy. We thank Drs Anne O’Garra, Gil Castro, Fabiani Frantz and Tiago Beites for critically reading the manuscript and Dr. Zewen Kelvin Tuong for insightful discussions.

