## Supplementary material for "Activation of nuclear receptors correlates with tuberculosis severity and is a target for host-directed therapy": Data Sheet 1.PDF

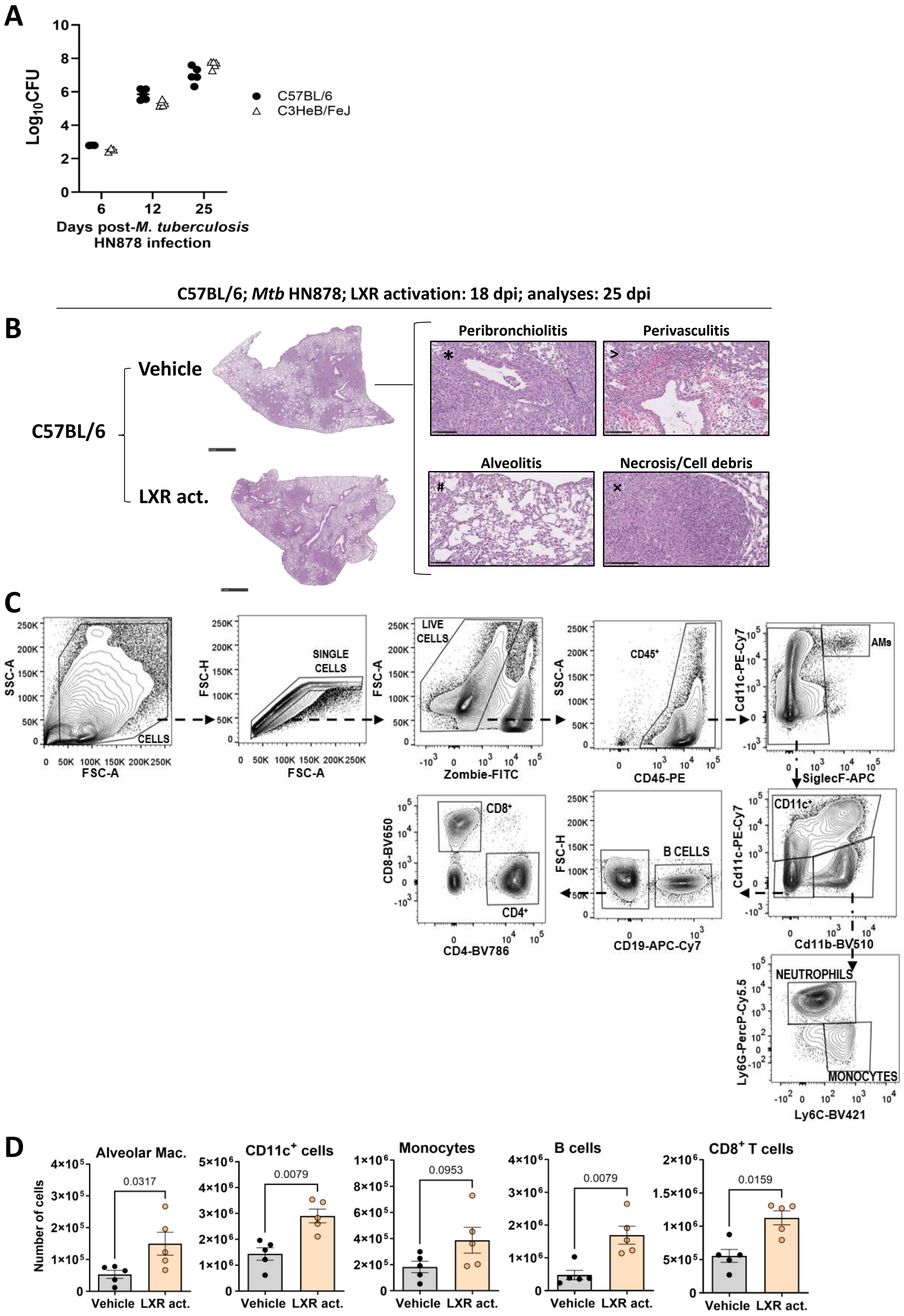


Supp. Fig. 1 (legend in the next page)

**Supp. Fig. 1. Activation of the LXR pathway during *M. tuberculosis* infection of C57BL/6 mice.** (A) Lung bacterial loads of C57BL/6 (circles) or C3HeB/FeJ (triangles) mice upon aerosol infection with *M. tuberculosis* HN878 (580 and 209 CFU delivered to the lung of C57BL/6 or C3HeB/FeJ mice respectively). Bacterial loads were determined on days 6, 12 and 25 post-infection. Each dot represents the mean ± SEM for 3-5 animals per time point in one experiment. (B) Representative images of hematoxylin-eosin stained lung sections for control (vehicle) or LXR agonist administered mice, starting on day 18 and analysed on day 25 post-infection. The images were used for the morphometric analyses presented in Fig. 3C. Scale bars 1 mm. Close ups show images representing the histologic parameters (peribronchiolitis, perivasculitis, alveolitis and necrosis/cell debris) used to obtain the combined lesion score presented in Fig. 3D. Scale bars 100 μm. (C) Gating strategy used to analyse the lung immune cell populations. (D) Cell numbers of distinct lung immune cell populations obtained for control (vehicle, grey) or LXR treated (orange) C57BL/6 mice. The LXR agonist was administered on day 18 and the lung immune composition analysed on day 25 post-infection. Data are mean ± SEM and each dot represents one mouse. Mann-Whitney U tests were used to identify statistical differences between the two groups. p-values are indicated and considered significant if ≤ 0.05.

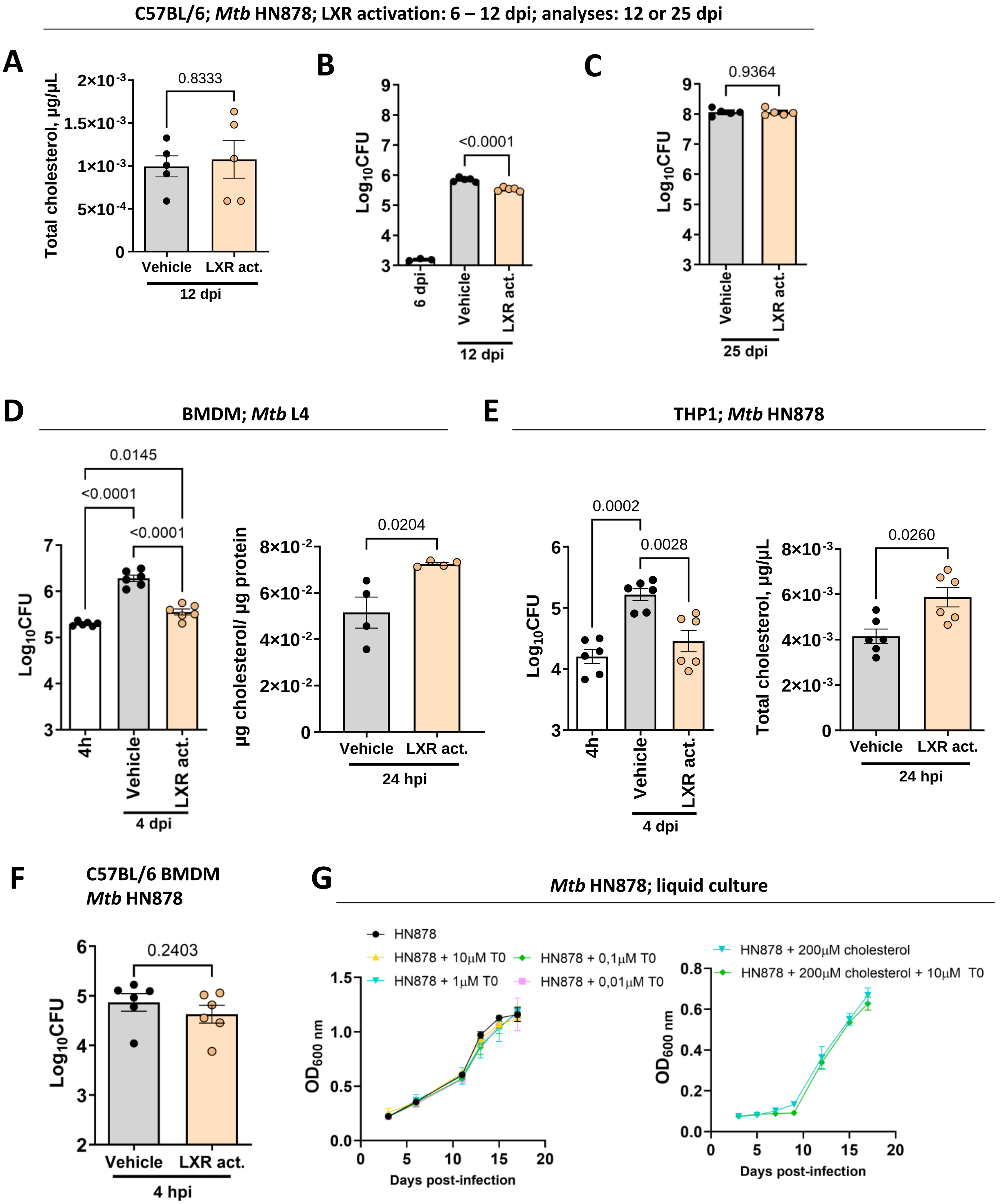


**Supp. Fig. 2. Effect of LXR activation in the lungs or BMDM of C57BL/6 mice infected with *M. tuberculosis*.** C57BL/6 mice were aerosol infected with *M. tuberculosis* HN878 (initial dose of 536 CFU delivered to the lung) and the LXR pathway activated between days 6 and 12 post-infection. On day 12 post-infection, the levels of lung cholesterol were measured in vehicle (grey bars) or LXR-treated (orange bars) mice (A). Lung bacterial burdens were assessed on days 6 (prior to LXR activation), 12 (after LXR activation) and 25 (experimental endpoint) post-infection (B, C). Bacterial loads and extracellular cholesterol levels detected in cultures of bone marrow-derived macrophages (BMDM) from C57BL/6 mice (D, F) or THP1 (E) cells infected with *M. tuberculosis* L4 (D) or HN878 (E, F). BMDM were infected at a MOI of 2 and THP1 cells at a MOI of 1. Where indicated, cell cultures were treated with 100 nM of the LXR activator. Bacterial loads were determined 4 h and 4 days post-infection. (A-F) Data are presented as mean ± SEM, with each dot representing an individual mouse or culture sample. Mann-Whitney U tests were used to identify statistical differences between the two groups. p-values are indicated and considered significant if ≤ 0.05. (G) Growth curves obtained for *M. tuberculosis* HN878 in axenic medium alone or with increasing concentrations of the LXR activator (left panel) or in media supplemented with 200 µM cholesterol in the absence or presence of the LXR activator (10 µM; right panel). dpi, days post infection; hpi, hours post-infection; T0, LXR activator

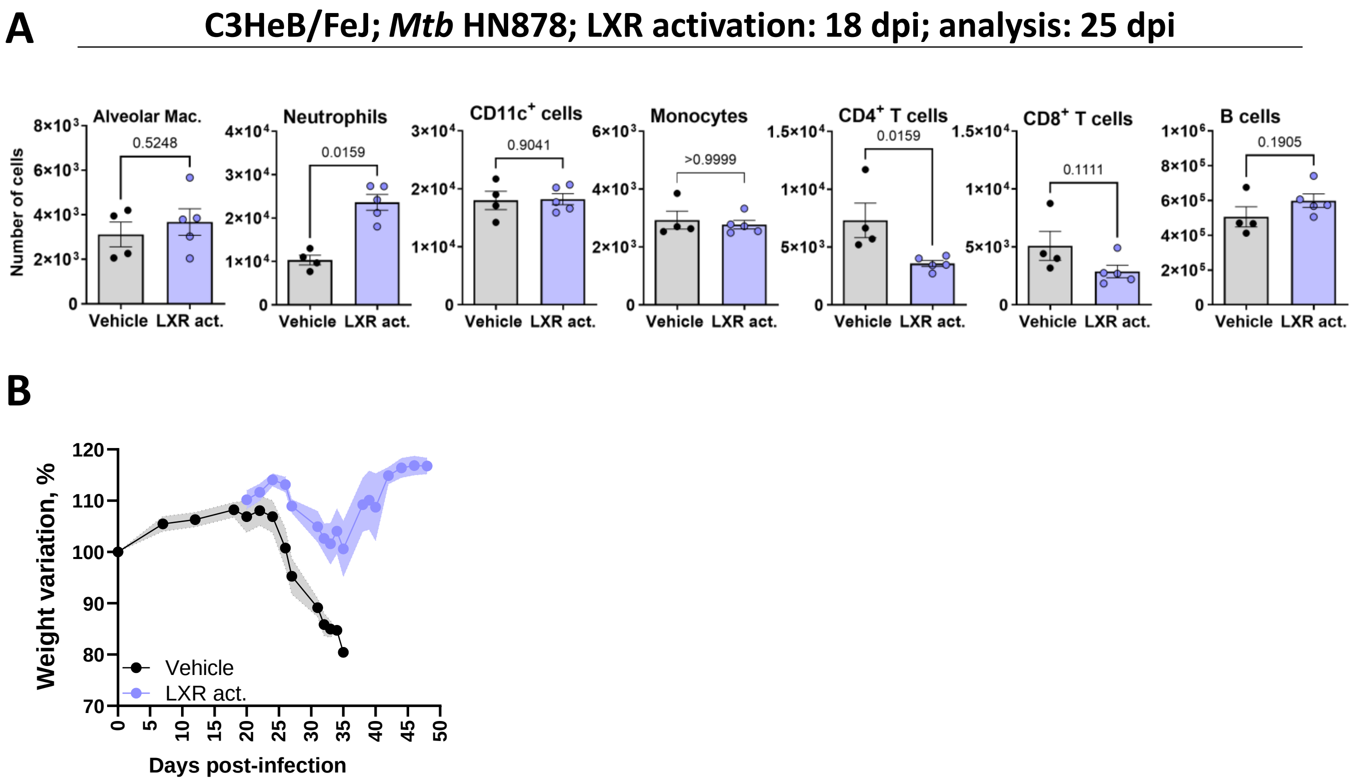


**Supp. Fig. 3. Lung immune cell populations and weight variation of control or LXR-treated *M. tuberculosis*-infected C3HeB/FeJ mice.** (A) C3HeB/FeJ mice were aerosol-infected with *M. tuberculosis* HN878 (average of 203 CFU delivered to the lung). On day 18 post-infection the LXR activator or the vehicle control were administered. The experiment was terminated on day 25 post-infection and the lung immune cell composition determined by flow cytometry for each experimental group. Gating strategies were as in Supp. Fig. 1C. Vehicle and treated groups are represented by grey or blue bars, respectively, and each dot represents a mouse. Data are mean ± SEM. Mann-Whitney U tests were used to identify statistical differences between the two groups. p-values are indicated and considered significant if ≤ 0.05. (B) Weight loss variation of control (vehicle; grey) or LXR-treated (blue) over time post-infection. The LXR agonist was administered from day 18 post-infection. The curves represent 2 independent experiments (average of 203 and 271 CFU delivered to the lung) with a total of 15 mice per group, pooled together, with SEM shaded.
