## Supplementary material for "Activation of nuclear receptors correlates with tuberculosis severity and is a target for host-directed therapy": Data Sheet 2.PDF

**Supplementary Table 3. List of primer-probes used in this study.**

| Gene | Primer - probe |
|---|---|
| *Abca1* | Mm00442646_m1 |
| *Abcg1* | Mm00437390_m1 |
| *Lpcat3* | Mm00520147_m1 |
| *Srebf1* | Mm00550338_m1 |
| *Arg1* | Mm00475988_m1 |
| *Arg2* | Mm00477592_m1 |
| *Nr1h3* | Mm00443451_m1 |
| *Apoe* | Mm01307193_g1 |
| *Hprt1* | Mm03024075_m1 |
| *Hmbs* | Mm01143545_m1 |

**Supplementary Table 4. List of flow cytometry antibodies used in this study.**

| Target | Clone | Fluorochrome | Catalogue* # | RRID |
|---|---|---|---|---|
| CD4 | GK1.5 | BV786 | 100453 | AB_2565843 |
| CD8 | 53-6.7 | BV650 | 100741 | AB_2563056 |
| CD11b | M1/70 | BV510 | 101245 | AB_2561390 |
| CD11c | N418 | PE-Cy7 | 117318 | AB_493568 |
| CD19 | 6D5 | APC-Cy7 | 115530 | AB_830707 |
| CD45 | 30-F11 | PE | 103106 | AB_312971 |
| Ly6C | HK1.4 | BV421 | 128014 | AB_1732079 |
| Ly6G | 1A8 | PercP-Cy5.5 | 127616 | AB_1877271 |
| SiglecF | S17007L | APC | 155508 | AB_2750237 |

* all Biolegend
